# Lipobiotin-capture magnetic bead assay for isolation, enrichment and detection of *Mycobacterium tuberculosis* from saliva

**DOI:** 10.1101/2022.03.04.483051

**Authors:** Julia Hansen, Katharina Kolbe, Inke R. König, Regina Scherliess, Marie Hellfritzsch, Sven Malm, Julia Zallet, Doris Hillemann, Karl-Heinz Wiesmüller, Christian Herzmann, Julius Brandenburg, Norbert Reiling

**Author notes:** Karlsruhe Institute of Technology, Eggenstein-Leopoldshafen, Germany (Sven Malm). equal contributing authors (Julius Brandenburg, Norbert Reiling).

## Abstract

**Background:** Pulmonary Tuberculosis (TB) is diagnosed through sputum samples. As sputum sampling is challenging in children and cachexic patients, the development of diagnostic tests using saliva appears promising but has been discouraged due to low bacterial load and the poor sensitivity. We present a novel and rapid method to enrich *Mycobacterium tuberculosis* (Mtb) from saliva, which may serve as a basis for a diagnostic saliva test.

**Methods:** **L**ipobiotin-functionalized **m**agnetic **b**eads (LMBs) were incubated with Mtb-spiked PBS and saliva from healthy donors as well as with saliva from TB patients. Flow cytometry was used to evaluate the capacity of the beads to bind Mtb, while real-time quantitative polymerase chain reaction (qPCR) was utilized to detect Mtb and estimate the amount of mycobacterial DNA in different sample types.

**Results:** We found that LMBs bind Mtb efficiently when compared to non-functionalized beads. The development of an qPCR assay based on the use of LMBs (LMB assay) allowed us to enrich mycobacterial DNA in spiked sample types, including PBS and saliva from healthy donors (enrichment of up to 8.7 fold). In Mtb-spiked saliva samples, we found that the LMB assay improved the detection rate of 10^2^ bacteria in a volume of 5 ml from 0 out of 15 (0%) to 6 out of 15 (40%). Consistent with that, the LMB assay increased the rate of correctly identified saliva samples from TB patients in two independent cohorts.

**Conclusions:** Implementation of the principle of the LMB-based assay may improve the sensitivity of existing diagnostic techniques, e.g. by functionalizing materials that facilitate Mtb sampling from the oral cavity.

## Introduction

Worldwide, tuberculosis (TB) is one of the top-ten causes of death and the leading cause of death from a single bacterial agent (1). Every year, one million children worldwide fall ill with TB, which corresponds to about 10% of all TB cases. Due to its early manifestation without a prolonged phase of latency, childhood TB is a key prognostic factor for the rate of infection in a population. However, children often develop no or non-specific symptoms and the established diagnostic tests are not always applicable to children, because children have difficulties in producing sputum (2). Furthermore, handicapped or weak patients are often unable to cough up enough material for analysis. About 30% of all patients are not able to produce sputum (3).

In pulmonary TB, bacteria are primarily found in the lungs but enter the pharynx and oral cavity through coughing. Therefore, saliva has long been considered a feasible and readily available diagnostic material. As early as 1909, Neild et al. detected mycobacteria in saliva with a Ziehl-Neelsen stain (4). Over the following decades, it was commonly accepted that saliva is not a suitable biomaterial because it contains only 0.1-1% of the bacterial concentration in sputum (5–8). However, PCR-based approaches were able to detect *Mycobacterium tuberculosis* (Mtb) in saliva, suggesting that saliva is a reasonable and valuable alternative when sputum could not be generated (9–11). Nevertheless, the low bacterial count in saliva remains a problem, because of reduced sensitivity (8). This could be changed through an assay enriching the bacteria from saliva prior to detection.

In this study, we present a novel method based on Lipobiotin-functionalized magnetic beads that can effectively bind and enrich Mtb from solutions and saliva for further processing with commercial PCR kits (5).

## Material and Methods

### Bacteria and colony forming units (CFU) analysis

mCherry10-expressing *Mycobacterium tuberculosis* (Mtb) was generated as described elsewhere (12). Mtb H37Rv was obtained from American Type Culture Collection ((ATCC) 27294, Manassas, VA, USA). Strains were cultured as described in detail elsewhere (13). In order to generate frozen stocks, mid-log-phase bacterial suspensions (OD_600_, 0.2 to 0.4) were stored at −80°C. After four weeks of storage, the number of bacteria (colony forming units, CFU) was determined by plating serial tenfold dilutions on 7H10 plates (0.05 % Tween 80 / 10 % heat-inactivated bovine serum in PBS) and counting colonies after 3-4 weeks of incubation at 37 °C.

For spiking experiments, frozen stocks of Mtb H37Rv were thawed, centrifuged (4000*×*g), bacteria re-suspended in PBS and a defined amount of this solution added to buffer or saliva samples. To address the impact of proteinase K plus DTT treatment on bacterial viability, Mtb H37Rv spiked PBS samples were incubated in the presence or absence of Proteinase K (18U/ml, Carl Roth, Karlsruhe, Germany) and Dithiothreitol (DTT, 2mM, Sigma-Aldrich, St. Louis, USA) for 60 minutes at 37°C and subsequently subjected to CFU analysis as described earlier.

### Lipobiotin-functionalization of magnetic beads

5.6 × 10^8^ magnetic beads (Micromer-M, polystyrene body, surface: PEG-COOH, 5μm; in dH_2_0) were incubated with 12.8 mg N-hydroxysuccinimide (NHS) and 6.4 mg 1-ethyl-3-(3-dimethylaminopropyl) carbodiimide (EDC) in 20% [v/v] 2-(N-morpholino)ethanesulfonic acid (MES) buffer (pH 6.3) and agitated for 1.5 hours. Activated beads were washed twice with PBS and then incubated with the biotinylated lipopeptide Lipobiotin (PHCKKKKK(Aca-Aca-Biotin) x 3 TFA, N-Palmitoyl-S-(1,2-bishexadecyloxy-carbonyl) ethyl-[R]-cysteinyl-[S]-lysyl-[S]-lysyl-[S]-lysyl-[S]-lysyl-[S]-lysine(ε-aminocaproyl-∈-aminocaproyl-biotinyl) x 3 CF3COOH (EMC Microcollections, Tübingen, Germany) at 1.5 mg/ml in carbonate buffer (pH 9.2) at RT for 4.5 hours. Functionalized beads were washed twice with PBS and incubated in 10mM ethanolamine (in carbonate buffer; pH 9.2) to block unreacted ester groups (1h, RT). Lipobiotin-functionalized magnetic beads (LMB) were washed, re-suspended in PBS and kept at 4°C. Non-functionalized magnetic beads (NMB) that were equally washed and re-suspended in PBS served as a control.

### Treatment of Mtb-spiked buffer samples with magnetic beads

Phosphate buffered saline (PBS) (0.5 ml) was spiked with Mtb H37Rv at the indicated concentrations. Subsequently, 4.6×10^6^ LMBs or NMBs were added and samples incubated at 37 °C for 1 hour on a thermomixer (60 rpm; Eppendorf, Hamburg, Germany). After this step, beads were retained in the tube by use of a magnet (DynaMag-2 Magnet, Life technologies) and washed two times with PBS. Finally, beads were re-suspended in Tris and EDTA (TE) buffer with pH 10 before proceeding further.

### Saliva collection

Saliva samples were collected without stimulation from healthy donors or from patients with drug-sensitive, sputum smear positive pulmonary TB at different times of the day and were subsequently stored at −80°C. Samples were pseudonymized and processed in an observer-blinded fashion. The study was approved by the Ethics Committee of the University of Luebeck (ref. 14-079A).

### Treatment of saliva samples with LMBs

The treatment with LMBs, also referred to as “LMB assay”, for enrichment and quantification of mycobacterial DNA in saliva samples can be summarized as following. Saliva samples were thawed and either spiked with Mtb H37Rv or directly pre-treated with Proteinase K (18U/ml) and Dithiothreitol (DTT, 2mM) for 45 minutes at 37°C (Figure 6, step 1). Subsequently, samples were centrifuged (150×g), and supernatants transferred into fresh tubes. 4.6×10^6^ LMBs were added and samples incubated for 60 minutes at 37°C (Step 2). Samples were centrifuged (3629×g), and beads retained in the tube by use of a magnet (DynaMag-2 Magnet, Life technologies) (Step 3). Subsequently, beads were washed twice with PBS and re-suspended in Tris and EDTA (TE) buffer (pH 10), and heat-treated for the release of the DNA (95°C, 30 mins). After purification and concentration by ethanol precipitation (step 4, for further details see below), samples were subjected to real-time quantitative PCR analysis (step 5, for details see below).

**Figure 1:**
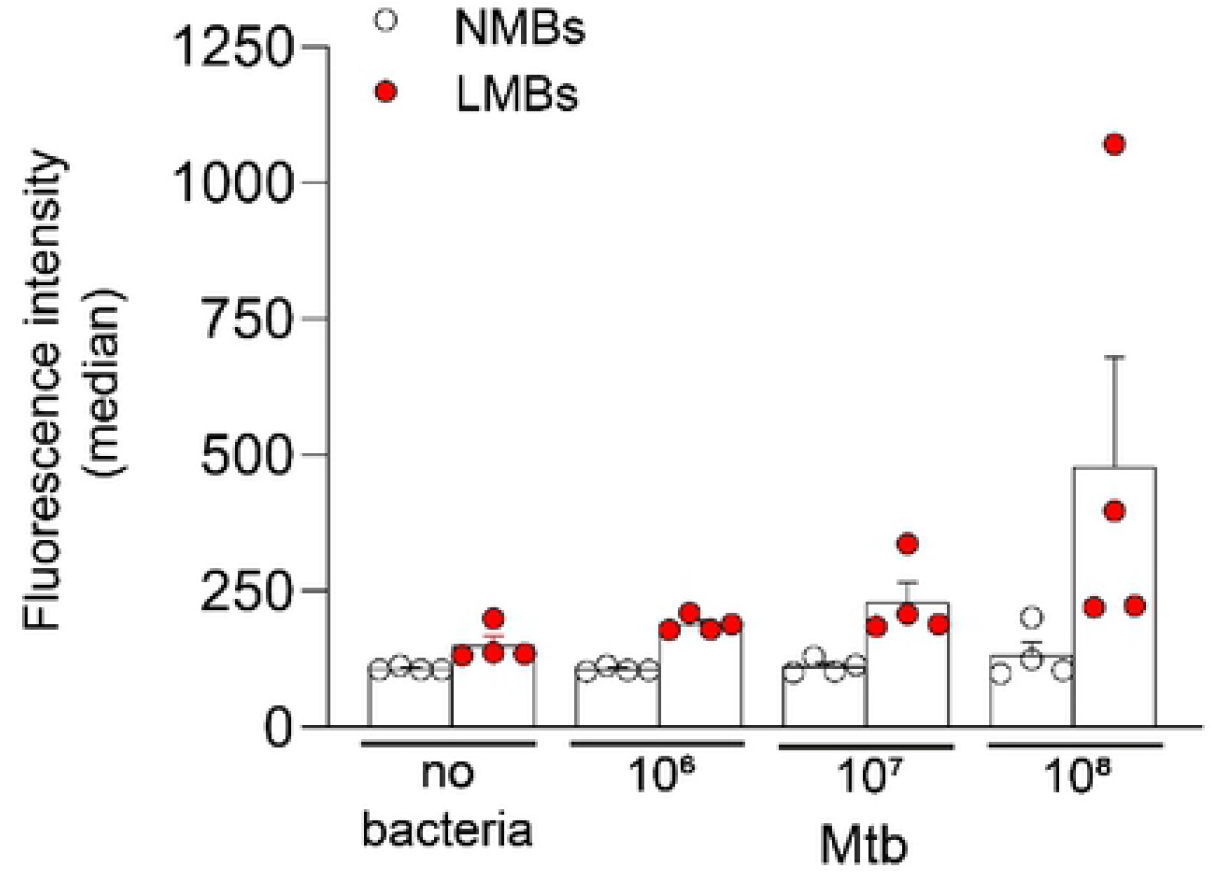
Mtb binds to Lipobiotin-functionalized magnetic beads (LMBs). mCherry-expressing Mtb bacteria were incubated in the presence of non-functionalized magnetic beads (NMBs) and lipobiotin-functionalized magnetic beads (LMBs) for 1 h at 37°C. Subsequently, beads were washed and subjected to flow cytometry analysis. The mean +/−SEM of the median fluorescence intensity of 4 independent experiments is shown. A non-parametric analysis of variance for repeated measurements was used as a statistical test, which included data from all doses of bacteria (NMBs vs. LMBs; p= 9.3 × 10^−27^).

**Figure 2:**
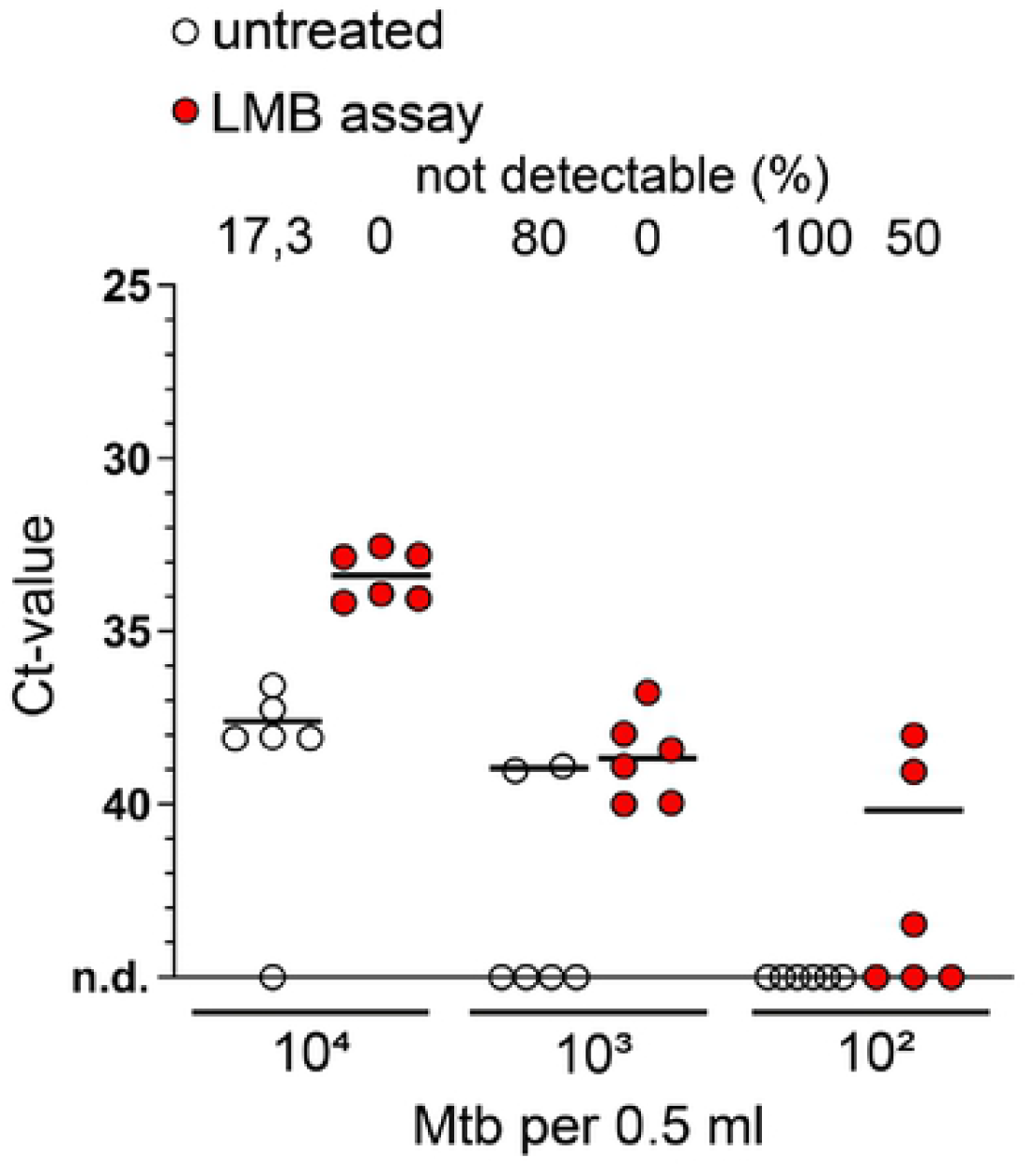
The effect of the LMB-assay on qPCR-based detection of Mtb in buffer. PBS samples were spiked with Mtb H37Rv and either left untreated or underwent pre-treatment with LMBs (LMB assay). The same volume of untreated or LMB pre-treated sample was subjected to heat treatment, DNA purification and qPCR analysis. Line at each condition depicts the mean of data from one representative out of two conducted experiments with six technical replicates. Above each condition, the percentage (%) of samples in which no signal could be detected (not detectable, n.d.) is given.

**Figure 3:**
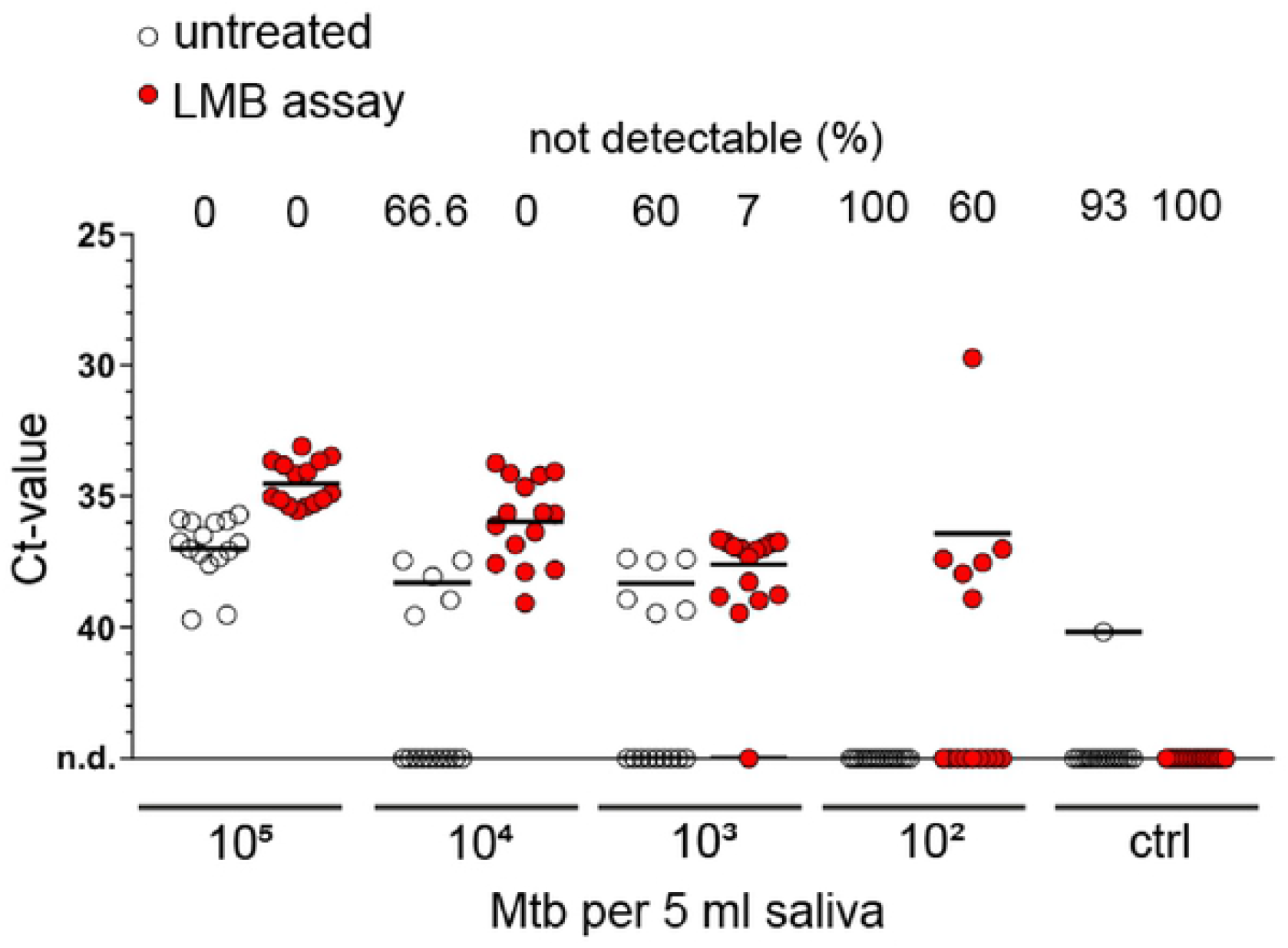
The effect of LMB assay on qPCR-based detection of Mtb in saliva. Saliva samples from healthy subjects were spiked with Mtb H37Rv and either left untreated or underwent pre-treatment with LMBs (LMB assay). The same volume of untreated or LMB pre-treated sample was subjected to heat treatment, DNA purification and qPCR analysis. Line at each condition indicates the mean of data from five independent experiments each consisting of three technical replicates. Above each condition, the percentage (%) of samples in which no signal could be detected (not detectable, n.d.) is given. Ctrl, control, no bacteria added. A non-parametric analysis of variance for repeated measurements was used as a statistical test, which included data from all doses of bacteria (untreated vs LMB assay; p=1.2 × 10^−35^).

**Figure 4:**
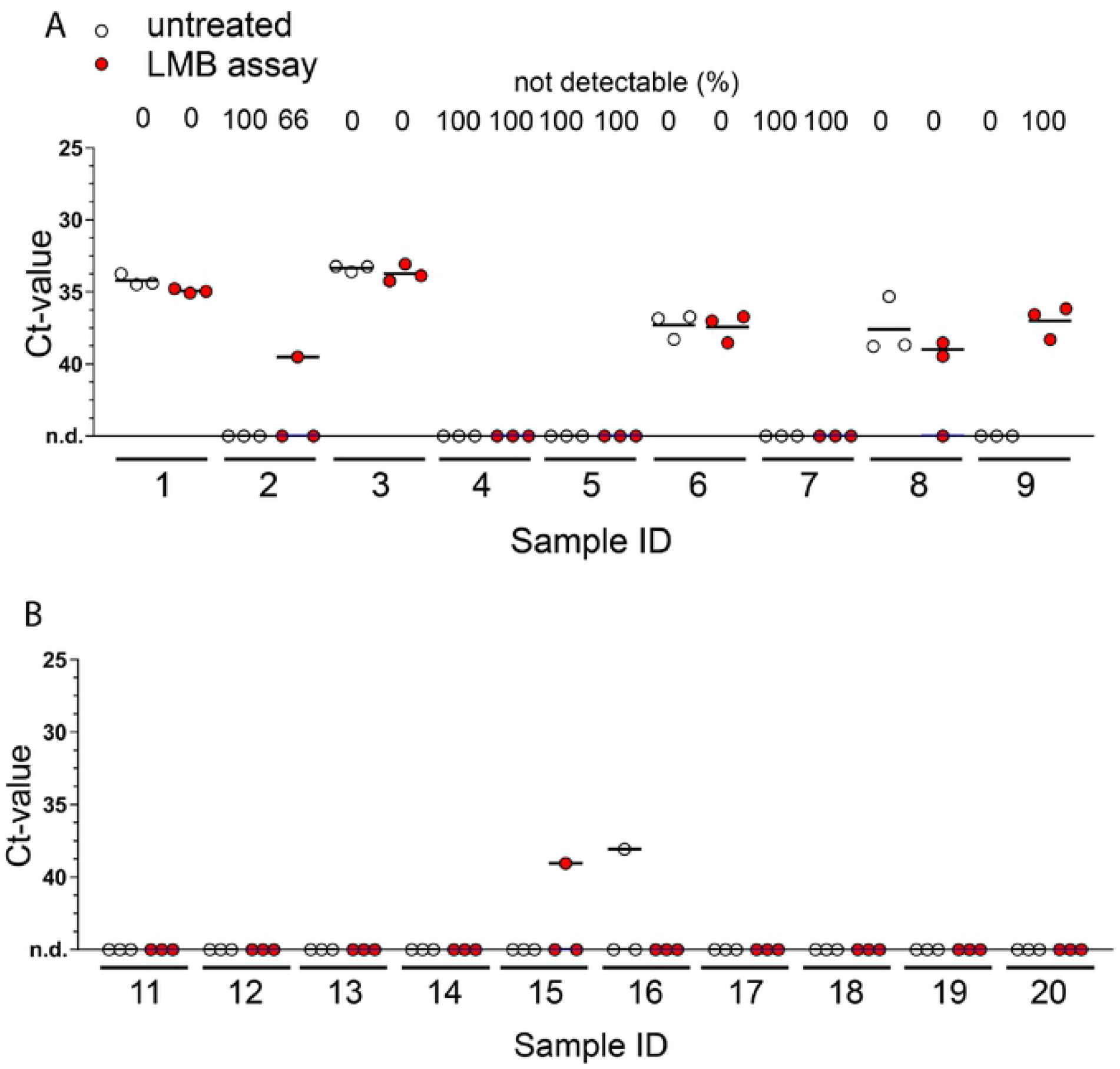
The effect of LMB assay on qPCR-based detection of Mtb in saliva from TB patients and healthy subjects (cohort #1). Saliva was collected from TB patients (A) and healthy subjects (B). Samples were divided into two fractions that were either left untreated or were incubated with LMBs (LMB assay) before undergoing heat treatment, DNA isolation, and qPCR analysis. Analysis of three technical replicates per condition is shown with the line at mean. For TB patients (A), the percentage (%) of samples in which no signal could be detected (not detectable, n d.) is given above the graph. Statistical analysis was conducted with a non-parametric analysis of variance for repeated measurements comparing untreated vs LMB assay-treated samples (A): p=0.631.

**Figure 5:**
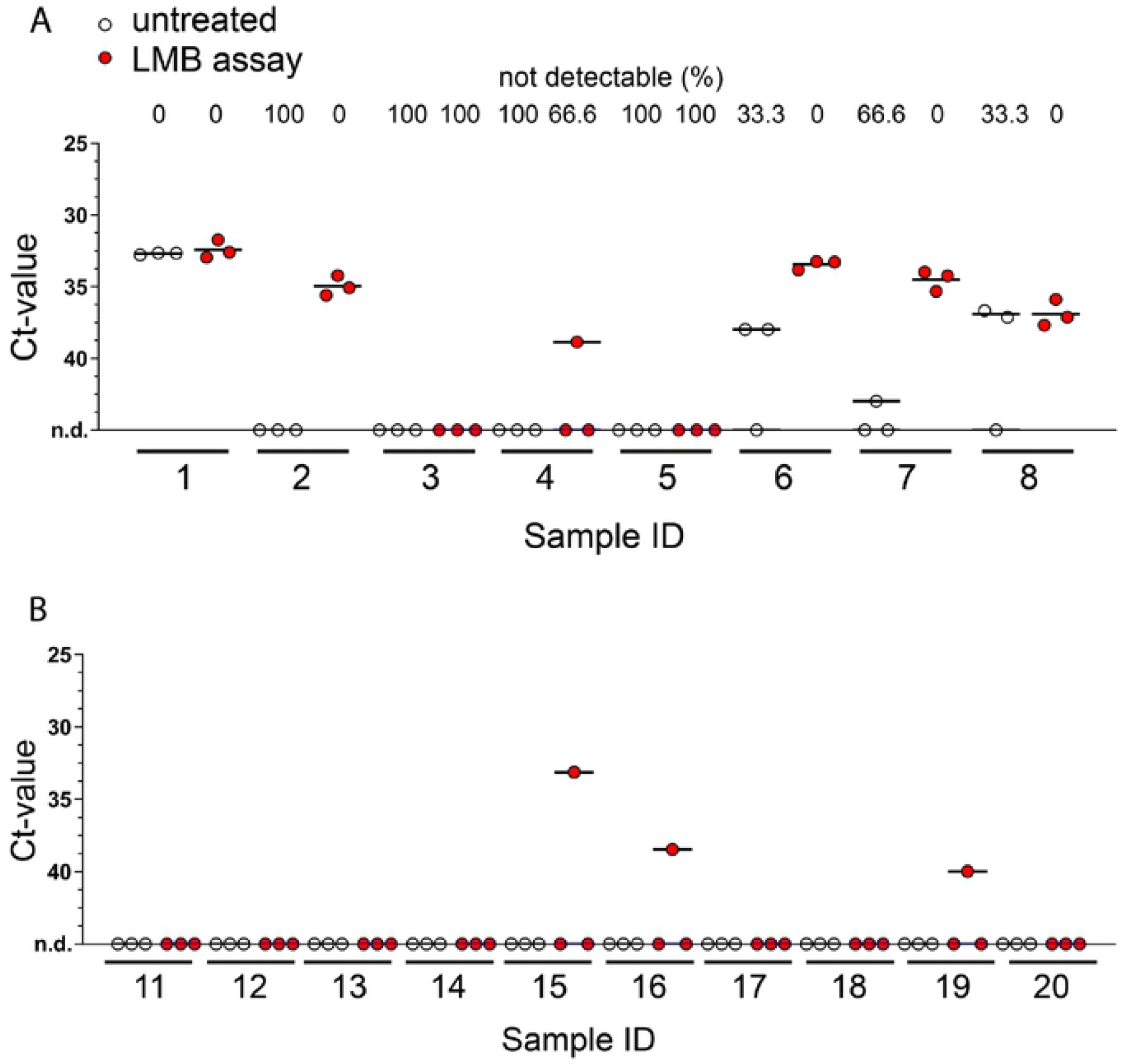
The effect of LMB assay on qPCR-based detection of Mtb in saliva from TB patients and healthy subjects (cohort #2). Saliva samples from TB patients (A) and healthy subjects (B) were divided into two fractions that were either left untreated or were incubated with LMBs (LMB assay) before undergoing heat treatment, DNA isolation, and qPCR analysis. Analysis of three technical replicates is shown with the line at mean +/−SEM. For TB patients (A), the percentage (%) of samples in which no signal could be detected (not detectable, n.d.) is given above the graph. Statistical analysis was conducted with a non-parametric analysis of variance for repeated measurements comparing untreated vs LMB assay-treated samples (A): p=0.0085).

**Figure 6:**
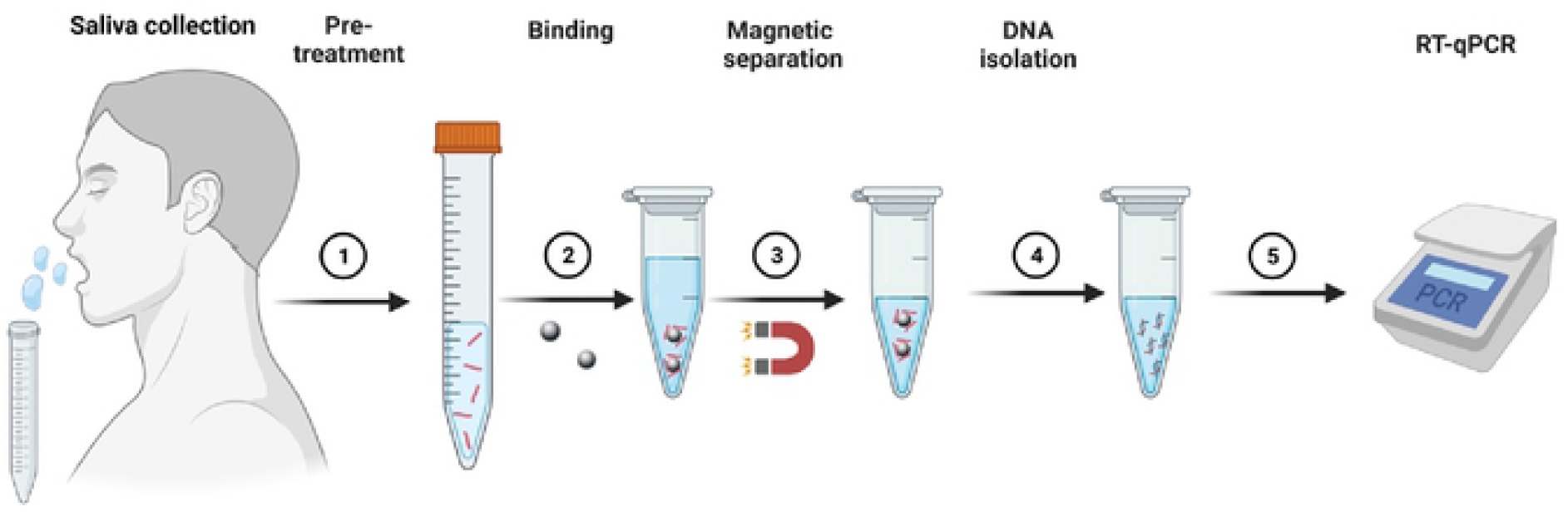
Schematic illustration of the Lipobiotin magnetic bead assay (LMB assay) After saliva collection, saliva samples were pre-treated with proteinase K (18U/ml) and dithiothreitol (DTT, 2mM) for 45 mins at 37°C (Step 1). Lipobiotin-functionalized magnetic beads (4.6×10^6^ LMBs) were added and samples incubated for 60 mins at 37°C (step 2). Subsequently, beads were isolated by centrifugation and byuse of a magnet (DynaMag-2 Magnet) (Step 3). After this step, samples were heat-treated (95°C, 30 mins) and extracted DNA purified and concentrated by ethanol precipitation (Step 4). Finally, samples were subjected to real-time quantitative PCR analysis (5).

### Rheology

Saliva samples were mixed with proteinase K (18U/ml) and DTT (2mM) or with the same volume of dH_2_O as control and incubated (45 minutes, 37°C, 60rpm) on a roller mixer. The rheological properties of the saliva were measured using a Bohlin Rheometer (Bohlin CVO 120 High Resolution, Malvern Instruments Ltd., UK) and the 40mm diameter plate-plate apparatus. The slit width was set to 150μm and the temperature was constant at 36°C. The change in viscosity of 1 ml of each sample was recorded with a shear profile of 1 s^−1^ to 100 s^−1^. For comparison the results were determined three times and of the profiles from 90 s^−1^ to 100 s^−1^ were used.

### Flow cytometry

In order to assess the binding of Mtb bacteria to beads, mCherry10-expressing Mtb (at the concentration indicated) in PBS were incubated with 1.57×10^6^ LMBs or UMBs for 60 mins at 37°C and beads were separated by use of a magnet (see above). Subsequently, beads were washed twice with PBS and subjected to flow cytometry analysis at a BD FACSCanto™ II (B&D Biosciences, Franklin Lakes, USA). Data was analyzed by use of FCS Express 7.10.0007 (DeNovo Software, Los Angeles, USA) or earlier versions.

### DNA extraction and purification by ethanol precipitation

Samples containing Mtb were heat-treated on a thermomixer (95°C, 30 minutes, 1400 rpm). Subsequently, in order to concentrate and purify the extracted DNA, samples were vortexed in the presence of ethanol (81% [v/v], VWR International, Radnor, USA) ammonium acetate (9% [v/v], Thermo Fisher Scientific, Waltham, USA) and 60 μg glycogen (Roche, Basel, Switzerland) and incubated overnight at −20°C. Following centrifugation (30 minutes, 16.200×g), the supernatant was discarded. This step was repeated. Subsequently, pellet was washed with ethanol (75% [v/v] and sample centrifuged (10 minutes, 16.200×g). Finally, the pellet was air dried for 15 minutes and dissolved in 5μl H_2_O before proceeding further.

### Real-time quantitative PCR (qPCR)

For detection of Mtb DNA, real-time quantitative PCR (qPCR) was performed based on a previously published protocol that detects a region in Rv2819 of Mtb (14). In a final volume of 10 μl TaqMan Universal PCR Master Mix (Roche), forward and reverse primer (nBjF(5’-aagcattcccttgacagtcgaa); nBjR(5’-ggcgcatgactcgaaagaag) 500nM each), fluorescein-(FAM)-BlackBerry-(BBQ)-non-Beijing-Probe-(5’-6FAM-tcatcaaagaccctcttggaaggccc-BBQ; 200nM), double distilled water and the isolated Mtb DNA were mixed in a 96 well plate. The plate was sealed and centrifuged (2 minutes, 180×g, 4°C) before analysis with the LightCycler 480 Instrument II (Roche; Preincubation, 95°C, 10 min; Denaturation, 92°C, 15s; Extension, 60°C, 60s; Cooling, 40°C, 10s for 50 cycles).

### Statistical Analysis

Statistical evaluation was performed by use of R 3.5.0 and the R package nparLD (Open Source; (15)). The data was analysed by use of non-parametric analyses of variance for repeated measurements (15) and the obtained p-value(s) were corrected for the number of tests performed. Data from all bacterial doses were included and, except for data depicted in figure 1, the statistical evaluation between untreated and LMB assay treated samples reported.

## Results

### Mtb binds to Lipobiotin-functionalized magnetic beads (LMBs)

To assess whether lipobiotin-functionalized magnetic beads (LMBs) are of potential use to enrich Mtb bacteria from patient material such as saliva, we assessed whether Mtb bacteria bind to LMBs and compared the binding to non-functionalized magnetic beads (NMBs). Thus, PBS-samples (0.5ml) were spiked with various numbers of fluorescent, mCherry-expressing Mtb bacteria (10^6^, 10^7^ or 10^8^; equivalent of 2×10^6^ - 2×10^8^ bacteria per ml) and incubated in the presence of LMBs or NMBs. Subsequently, magnetic beads were isolated, washed and subjected to flow cytometry analyses. As shown in Figure 1, the median fluorescence intensity increased in a dose-dependent manner with increasing numbers of added bacteria when samples were incubated with LMBs. In contrast, there was no substantial increase in median fluorescence intensity for samples incubated with NMBs, irrespectively of the amount of bacteria added. Considering data from all bacterial doses, a highly significant increase in fluorescence intensity was observed when LMBs were compared to NMBs (p= 9.3 × 10^−27^), strongly suggesting that Mtb bacteria bind to lipobiotin-functionalized beads, whereas only minimal or no binding occurs with non-functionalized beads.

### LMB pre-treatment (LMB assay) improves qPCR-based detection of mycobacterial DNA in Mtb-spiked buffer samples

qPCR is utilized as a molecular diagnostic test for tuberculosis by detecting mycobacterial DNA in samples. Thus, we assessed whether utilization of LMBs could improve qPCR-based detection of Mtb. In a first set of experiments, we spiked various amounts of Mtb H37Rv (10^2^, 10^3^ and 10^4^, equivalent of 2×10^2^-2×10^4^ bacteria per ml) to PBS samples (0.5 ml), which were then left untreated or underwent pre-treatment with LMBs. The magnetic beads were separated from the samples by use of a magnetic stand and – after two washing steps – re-suspended in PBS. The same volume of untreated or LMB pre-treated sample was subjected to heat treatment (95°C, 30 min), DNA purification and qPCR analysis detecting a well-established target in Mtb (14). As depicted in Figure 2, with 10^4^ bacteria, Mtb was not detectable in 1 out of 6 (17.3%) of the cases when samples were untreated, while all samples were tested positive when samples were pre-treated with LMBs (LMB assay). Notably, treatment reduced the average ct-values from 37.6 to 33.9 indicative of an increased concentration of Mtb DNA in samples treated with the LMB assay. After determining the reaction efficacy of the conducted qPCR (E= 1.77, compare Figure S2) we calculated the relative DNA concentration of the samples. For 10^4^ bacteria, this analysis revealed an ~8.4-fold enrichment of Mtb DNA in LMB-assay treated samples when compared to untreated samples. In line with this, with 10^3^ bacteria, Mtb was undetectable in 4 out of 6 (66.6%) of the cases when samples were untreated, while all samples were tested positive when samples were pre-treated with the LMB assay. Similarly, with 10^2^ bacteria, Mtb was not detected when samples were left untreated, but in 50% of the LMB-assay treated samples. In summary, LMB pre-treatment substantially reduced the mean ct-values and the rate of false negative PCR results. Results from this first set of experiments analysing aqueous, Mtb-spiked samples suggest that LMB pre-treatment has the potential to improve qPCR-based detection of Mtb also in other biological fluids.

### Treatment with Proteinase K plus DTT does not affect Mtb viability, but reduces viscosity of saliva samples

In contrast to aqueous solution, saliva has a higher viscosity, which may interfere with the binding of Mtb to the LMBs. To circumvent this issue, we aimed to use the protease proteinase K and the reducing agent dithiothreitol (DTT). This treatment, however, could affect viability of Mtb. Thus, we first incubated Mtb for 60 minutes at 37°C with Proteinase K (18U/ml) and DTT (2mM) and plated the bacteria on solid 7H10 agar, revealing similar CFU counts between untreated and Proteinase K/DTT samples (Figure S1). Second, to test the anti-viscosity effect of these compounds, we incubated saliva samples for 45 minutes at 37°C in the absence and presence of Proteinase K (18U/ml) plus DTT (2mM). Rheological analyses revealed that the viscosity of the saliva samples varied considerably between donors (Table 1). Importantly, treatment with Proteinase K plus DTT for already 45 minutes reduced the viscosity of saliva samples in all donors compared to the respective untreated controls. In summary, our data show that Proteinase K plus DTT treatment reduces viscosity of saliva samples, while not affecting viability of Mtb.

**Table 1:** Viscosity of saliva samples with and without Proteinase K / DTT treatment.

|  | Native viscosity,<br>(Pa·s·10<sup>-4</sup>) | Viscosity after proteinase K/<br>DTT treatment,<br>(Pa·s·10<sup>-4</sup>) | Viscosity<br>reduction<br>(%) |
| --- | --- | --- | --- |
| Sample 1 | 26.24 (±0.38) | 4.26 (±0.77) | 83.7 |
| Sample 2 | 16.18 (±0.48) | 3.85 (±0.29) | 76.2 |
| Sample 3 | 8.14 (±0.99) | 3.82 (±0.44) | 53.0 |
| Sample 4 | 7.19 (±0.12) | 3.97 (±0.32) | 44.7 |
| Sample 5 | 4.93 (± 0.72) | 3.86 (±0.59) | 21.7 |

### LMB assay improves detection of mycobacterial DNA in Mtb-spiked saliva samples

To test whether the LMB assay improves qPCR-based detection of Mtb in saliva, samples obtained from healthy donors (5 ml) were incubated with Proteinase K plus DTT, spiked with different amounts of Mtb (10^2^ – 10^5^ per 5ml; equivalent of 2×10^1^ - 2×10^4^ bacteria per ml) and were either left untreated or LMB-assay treated prior to heat-treatment, DNA extraction and qPCR analysis. With 10^5^ bacteria, all samples (15 in each group) were tested positive, irrespective of whether the LMB-assay was used or not (Figure 3). Importantly, the mean ct values decreased from 36.99 to 34.5, which represents an enrichment of DNA concentration by performing the LMB assay of 4.2-fold, when considering the reaction efficiency of the qPCR (E=1.77, see Figure S2). With 10^4^ bacteria, Mtb was not detectable in 10 out of 15 (66.6%) of the untreated samples, whereas detection rate remained at 100% with prior LMB-Assay. Similarly, with 10^3^ bacteria, Mtb was not detectable in 9 out of 15 (60%) of the cases, while the LMB assay allowed Mtb detection in 93% of the samples. Strikingly, the LMB-Assay allowed the detection of 10^2^ Mtb in 40% of the samples (6 out of 15), whereas Mtb DNA detection was not possible without prior LMB-Assay (0 out of 15). Importantly, when integrating data from all bacterial doses, ct-values were significantly reduced when statistically comparing LMB assay-treated with untreated samples (p=1.2 × 10^−35^). In summary, our data demonstrate that the LMB-Assay improves qPCR-based detection of Mtb in spiked saliva samples, suggesting it to be a valuable tool to enrich bacteria even from biological fluid with a high viscosity.

### Mtb DNA detection in saliva samples from TB-patients

To assess whether the LMB assay can improve a saliva-sample based diagnosis of pulmonary tuberculosis, two cohorts were tested in an observer-blinded manner, each consisting of ten saliva samples from healthy subjects (as a control for specificity) and eight and nine saliva samples from sputum-smear positive patients, respectively (Figure 4 and 5). One fraction of each sample was left untreated (control), while another fraction was incubated with LMBs before undergoing heat treatment, DNA isolation, and qPCR analysis (LMB assay; workflow as shown in Figure 6). Notably, from each saliva sample, three technical replicates were conducted and detection of Mtb was considered positive if any values were recorded positive. According to this, in the first cohort (Figure 4A), Mtb was detected in 44,4% of the TB patients when samples were left untreated (Sample ID 1, 3, 6 and 8; 4 out of 9), while detection of Mtb improved to 66,67% when LMB assay was employed prior to the qPCR (Sample ID 1, 2, 3, 6, 8, 9 and 10; 6 out of 9). However, the average ct-values in the four patients detected with both methods did not change, and there was no statistically significant difference between the groups (p=0.6079). In addition, in one patient (sample ID 9), qPCR detection after performing the LMB assay appeared less robust, as positive results were obtained in only two out of three cases in the technical replicates. Finally, there were no differences in the healthy subject’s samples with 10% false positive results for control and LMB assay (Figure 4B).

In the second cohort (Figure 5), positive qPCR signals were detected in 50% and 75% of cases, respectively, when untreated and LMB assay-treated samples were compared (Figure 5A). In two patients, Mtb was detected only when the LMB assay was applied (Sample ID 2 and 4). In four patients, both methods lead to a positive detection (Sample ID 1, 6, 7, 8), but the LMB assay showed a more reliable positive readout of the replicates and lowered the mean ct-value in two patients substantially (Sample ID 6 and 7). Consistent with that, we found significantly reduced ct-values in LMB-assay treated saliva samples when compared to untreated samples (p=0.0085), confirming that the LMB assay improves qPCR-based detection of mycobacterial DNA in this cohort. Surprisingly, the LMB assay led to three single positive PCR results (10% out of 30 tests performed) in saliva samples from healthy subjects compared to those that were left untreated (Figure 5, sample ID 15, 16 and 19). Consequently, we again collected saliva (2 of 3 donors were available), re-examined the samples and this time obtained negative qPCR results, regardless of whether the LMB test was performed or not (Figure S3, sample ID 15 and 19).

In summary, our data from two small cohorts show that more TB patients can be correctly identified from saliva analysis when samples were subjected to LMB assay prior to analysis.

## Discussion

In this study, we present a novel and rapid method to enrich *Mycobacterium tuberculosis* (Mtb) from saliva based on lipobiotin-functionalized magnetic beads (LMBs). The use of the LMB assay in samples spiked with Mtb increased the mycobacterial DNA concentration by 4.2-8.7-fold, thereby substantially improving the qPCR-based detection of Mtb. In saliva samples from healthy donors that were spiked with Mtb, we found that the LMB assay improves the detection rate of 10^2^ bacteria in a volume of 5 ml from 0 out of 15 (0%) to 6 out of 15 (40%). In addition, we show that more TB patients can be correctly identified from saliva by qPCR when samples were treated with the LMB assay. When combining data from two small patient cohorts, 8 out of 17 TB patients tested were correctly identified by qPCR when saliva was left untreated, while the detection rate increased to 12 out of 17 when saliva was treated with the LMB assay. Our findings may open a diagnostic door towards an improved Mtb detection even in patients who are unable to expectorate.

We provide evidence that the LMB assay improves qPCR-based detection of Mtb in spiked samples, but also in saliva derived from TB patients. The LMB assay is based on the interaction of Mtb with bead-bound lipobiotin, a molecule previously used to isolate Mtb-containing intracellular compartments from macrophages (16). During functionalization, lipobiotin is covalently bound to the polyethylene glycol (PEG) layer on the bead surface, leaving the lipophilic part of the molecule containing three fatty acids available for attachment of mycobacteria. In the first cohort from sputum smear-positive patients diagnosed with TB, the Mtb detection rate from saliva was slightly improved, but the LMB assay did not significantly enhance mycobacterial DNA concentration in the samples. In the second cohort investigated, the LMB-assay not only reduced the number of false negative results, but also enriched mycobacterial DNA in the samples significantly, supporting our pre-clinical findings from Mtb-spiked saliva samples. These discrepancies between the cohorts may have been due to the different bacterial loads of the patients but may also be a result of issues with the functionalization of the batch of beads used in cohort #1. Based on our observations, we hypothesized that the batch of beads used for cohort #1 may have had a lower capture molecule density on the bead surface, however we currently have no data to proof this hypothesis. Thus, the issue of consistent functionalization of the magnetic beads should be considered and addressed in future studies. A homogeneous surface coating would facilitate reliable Mtb detection from saliva, but may require additional technical development steps, which were beyond the focus of our study.

To prove the presence of Mtb bacteria in body fluids such as saliva, not only mycobacterial DNA but also mycobacterial proteins can be detected. Recently, it has been shown that the detection of the mycobacterial proteins MPT64 and CFP-10 by immuno-PCR can be improved by using functionalised gold nanoparticles. Similarly to the method we present here, this approach can improve the detection of Mtb in body fluids (17).

The developed LMB-Assay proved to be rather resilient to environmental factors, which is important for the implementation in high-prevalence countries with fewer resources. Natural variations of saliva viscosity were overcome by the pre-treatment with proteinase K and DTT, which did not negatively influence Mtb viability or detection. The principle of the LMB-Assay that we describe has the potential to improve TB diagnostics even in low resource settings. Currently, our investigations were limited to ex-vivo experiments on bacterial enrichment. Ideally, lipobiotin as a capture molecule would be attached to a chewable, porous matrix that could collect Mtb directly from the oral cavity. This diagnostic chewing gum or sponge would not only collect Mtb from saliva but also from dental and mucosal surfaces by abrasive shearing. Especially in children and patients that are unable to expectorate sputum, this approach is worth investigating. We are convinced that implementing the principle of the LMB-Assay into current ex-vivo diagnostic algorithms could improve the sensitivity of existing tests.

## Conclusion

An innovative LMB assay was developed to enrich Mtb from saliva samples. Lipobiotin coated beads interacting with the bacteria were able to bind Mtb in buffer and saliva, enrich their DNA and augment the detection by qPCR. In small cohorts of TB-patients, we showed that the LMB-Assay has a potential to improve the sensitivity of TB diagnostics. Eventually, using the ligand on a chewable matrix, our method may be suitable for the development of a diagnostic chewing gum for the detection of TB.

## References

1. Global tuberculosis report 2020 [Internet]. 2020 [cited 15.12.2021]. Available from: https://www.who.int/publications/i/item/9789240013131.

2. Diagnosis of childhood TB [Internet]. Available from: https://www.who.int/tb/challenges/ChildhhoodTB_section2.pdf.

3. Barnes P, Davies, P.D.O., & Gordon, S.B.. Clinical Tuberculosis (4th ed.). 2008.

4. Western GT. THE ROLE OF THE SALIVA IN THE TRANSMISSION OF TUBERCLE. The Lancet. 1909;173(4470):1276.

5. Kolbe K. New carbohydrate derivatives as tools to bind and metabolically label strains of the Mycobacterium tuberculosis complex. 2016. https://macau.uni-kiel.de/receive/diss_mods_00020638.

6. Yeager H, Jr., Lacy J, Smith LR, LeMaistre CA. Quantitative studies of mycobacterial populations in sputum and saliva. Am Rev Respir Dis. 1967;95(6):998–1004.

7. Holani AG, Ganvir SM, Shah NN, Bansode SC, Shende I, Jawade R, et al. Demonstration of mycobacterium tuberculosis in sputum and saliva smears of tuberculosis patients using ziehl neelsen and flurochrome staining-a comparative study. J Clin Diagn Res. 2014;8(7):Zc42–5.

8. Munot PP, Mhapuskar AA, Ganvir SM, Hazarey VK, Mhapuskar MA, Kulkarni D. Detection of Acid Fast Bacilli in Saliva using Papanicolaou Stain Induced Fluorescence Method Versus Fluorochrome Staining: An Evaluative Study. J Int Oral Health. 2015;7(7):115–20.

9. Gonzalez Mediero G, Vazquez Gallardo R, Perez Del Molino ML, Diz Dios P. Evaluation of two commercial nucleic acid amplification kits for detecting Mycobacterium tuberculosis in saliva samples. Oral Dis. 2015;21(4):451–5.

10. Wood RC, Luabeya AK, Weigel KM, Wilbur AK, Jones-Engel L, Hatherill M, et al. Detection of Mycobacterium tuberculosis DNA on the oral mucosa of tuberculosis patients. Scientific Reports. 2015;5(1):8668.

11. Mesman AW, Calderon R, Soto M, Coit J, Aliaga J, Mendoza M, et al. Mycobacterium tuberculosis detection from oral swabs with Xpert MTB/RIF ULTRA: a pilot study. BMC Research Notes. 2019;12(1).

12. Zelmer A, Carroll P, Andreu N, Hagens K, Mahlo J, Redinger N, et al. A new in vivo model to test anti-tuberculosis drugs using fluorescence imaging. Journal of Antimicrobial Chemotherapy. 2012;67(8):1948–60.

13. Reiling N, Homolka S, Walter K, Brandenburg J, Niwinski L, Ernst M, et al. Clade-Specific Virulence Patterns of Mycobacterium tuberculosis Complex Strains in Human Primary Macrophages and Aerogenically Infected Mice. mBio. 2013;4(4):e00250–13.

14. Hillemann D, Warren R, Kubica T, RüSch-Gerdes S, Niemann S. Rapid Detection of Mycobacterium tuberculosis Beijing Genotype Strains by Real-Time PCR. Journal of Clinical Microbiology. 2006;44(2):302–6.

15. Brunner E. Nonparametric analysis of longitudinal data in factorial experiments. In: Domhof S, Langer F, editors. New York, NY:: J. Wiley; 2002.

16. Steinhäuser C, Heigl U, Tchikov V, Schwarz J, Gutsmann T, Seeger K, et al. Lipid-labeling facilitates a novel magnetic isolation procedure to characterize pathogen-containing phagosomes. Traffic (Copenhagen, Denmark). 2013;14(3):321–36.

17. Dahiya B, Prasad T, Singh V, Khan A, Kamra E, Mor P, et al. Diagnosis of tuberculosis by nanoparticle-based immuno-PCR assay based on mycobacterial MPT64 and CFP-10 detection. Nanomedicine (Lond). 2020;15(26):2609–24.

